# Two distinct networks containing position-tolerant representations of actions in the human brain

**DOI:** 10.1101/2021.06.17.448825

**Authors:** Elahé Yargholi, Gholam-Ali Hossein-Zadeh, Maryam Vaziri-Pashkam

## Abstract

Humans can recognize other people’s actions in the social environment. This action recognition ability is rarely hindered by the movement of people in the environment. The neural basis of this tolerance to changes in the position of observed actions is not fully understood. Here, we aimed to identify brain regions capable of generalizing representations of actions across different positions and investigate the representational content of these regions. fMRI data were recorded from twenty-two subjects while they were watching video clips of ten different human actions in Point Light Display format. Each stimulus was presented in either the upper or the lower visual fields. Multivoxel pattern analysis and a searchlight technique were employed to identify brain regions that contain position-tolerant action representation: linear support vector machine classifiers were trained with fMRI patterns in response to stimuli presented in one position and tested with stimuli presented in another position. Results of this generalization test showed above-chance classification in the left and right lateral occipitotemporal cortex, right intraparietal sulcus, and right post-central gyrus. To explore the representational content of these regions, we constructed models based on the objective measures of movements and human subjective judgments about actions. We then evaluated the brain similarity matrix from the cross-position classification analysis based on these models. Results showed cross-position classifications in the lateral occipito-temporal ROIs were more strongly related to the subjective judgments, while those in the dorsal parietal ROIs were more strongly related to the objective movements. An ROI representational similarity analysis further confirmed the separation of the dorsal and lateral regions. These results provide evidence for two networks that contain abstract representations of human actions with distinct representational content.

## Introduction

Humans can rapidly and accurately recognize other people’s actions in the social environment. People’s movement in the environment introduces often dramatic changes in the viewpoint, size, and position of their image on the retina. This confounding variability renders the action recognition task computationally challenging. Yet, humans’ ability to recognize actions is rarely hindered by this variability. A brain region that subserves human action recognition ability is expected to: 1) contain information about various human actions and 2) contain action representations that are tolerant to the variability in the visual input. Here, using functional MRI and multi-voxel pattern analysis, we aim to identify such regions using changes in the position of the actor as the source of variability. After identifying these regions, we aim to explore their representational content and determine to what extent they relate to human subjective judgments about actions or objective measures of body movements.

Previous studies that have explored tolerance of visual representations to variability in the input have mostly focused on static pictures of objects and human bodies. Regions in both occipitotemporal (Konen and Kastner, 2008; Cichy et al., 2011, 2013; Anzellotti et al., 2013; Ramírez et al., 2014) and parietal cortices (Konen and Kastner, 2008) have been identified that contain position, size, and viewpoint tolerant representation of static objects (Konen and Kastner, 2008; Mur et al., 2010; Cichy et al., 2011, 2013; Anzellotti et al., 2013; Ramírez et al., 2014; Xu and Vaziri-Pashkam, 2020). This characteristic has been proposed as a defining feature of regions that contribute to human abstract object knowledge (Dicarlo and Cox, 2007). In the domain of observed actions, despite the large body of literature that identifies regions in the occipitotemporal and parietal cortices that respond to (Casper et al., 2010; Kalenine et al., 2010; Grosbras et al., 2012; Watson et al., 2013; Urgesi et al., 2014) and contain information about (e.g., Wheaton et al., 2004; Jastorff et al., 2010; Abdollahi et al., 2013; Lignau and Downing, 2015; Ferri et al., 2015; Hafri et al., 2017; Wurm et al., 2017b; Urgen et al., 2019; Tucciarelli et al., 2019; Tarhan and Konkle, 2020; Urgen and Orban, 2021) observed actions, evidence for the tolerance of these representations to changes in the position of the actors is still scant.

Previous studies have extensively investigated the tolerance of action representations to changes in viewpoint in both monkeys (Oram and Perrett, 1996; Vangeneugden et al., 2014; Caggiano et al., 2011; Maeda et al., 2015; Maranesi et al., 2015; Barz et al., 2017; Simone et al., 2017; Livi et al., 2019; Albertini et al., 2020; Lanzilotto et al., 2020) and humans (Grossman et al., 2010; Ogawa and Inui, 2011; Oosterhof et al., 2012; Tucciarelli et al., 2015; Isik et al., 2016). Results of these human studies provide evidence for viewpoint tolerance in the lateral occipito-temporal (Grossman et al., 2010; Tucciarelli et al., 2015) and parietal cortices (Ogawa and Inui, 2011; Oosterhof et al., 2012). In contrast to viewpoint tolerance, position tolerance has not been extensively explored. Grossman et al. (2010) employed an fMRI adaptation approach to investigate position tolerance in the human brain. They found that a region in human STS shows tolerance to variations in the position of observed actions. They did not systematically investigate the position sensitivity of parietal action selective regions. The fMRI adaptation technique used in this study may not have sufficient power in identifying the full scope of position tolerance in action selective regions compared to other techniques such as multi-voxel pattern analysis (MVPA, Haxby, 2012) that directly investigate the representational content of a region. Roth and Zohary (2015) employed MVPA (correlation and cross-decoding) to study position tolerance during observation of tool-grasping movements in humans. Their stimuli were presented in various locations, and position tolerance was examined for presentations in the left and right visual fields. They discovered that position information is gradually lost, and hand/tool identity information is enhanced along the posterior-anterior axis in the dorsal stream. However, their study was limited to grasping movements. It is not obvious if their results would generalize to full-body movements.

Other studies that have employed MVPA have often used natural stimuli in which actions happen at different positions, but they have averaged the responses across stimuli instead of systematically investigating the tolerance of representations across changes in position. Therefore it is hard to speculate about the extent of position tolerance in action selective regions from these studies (Hafri et al., 2017; Wurm et al., 2017b; Tarhan and Konkle, 2020). As such, a systematic investigation of position tolerance across dorsal and ventral action selective regions is still missing in the literature. In this study, we will use MVPA and cross-position decoding using a support vector machine classifier to search for position tolerant representations of actions in the human brain. Finding these regions, we will then investigate their representational content using a representational similarity analysis (Kriegeskorte et al., 2008).

Several theories have been proposed to characterize the role of individual action selective regions in the processing of action stimuli. In the occipito-temporal cortex, a division of labor has been suggested between regions such as EBA that process the form of the body and regions such as STS that process the movement of the body (Giese & Poggio, 2003; Peelen et al., 2006; Grossman et al., 2010; Michels et al., 2005; Downing et al., 2006; Vangeneugden et al., 2016 refs from Vangeneugden; Jastorff and Orban, 2009). These studies do not elaborate on the representational content of these regions. A few studies have taken a step further to establish the principles of coding in action selective regions. Within the lateral occipital cortex, Wurm et al. (2017b) have suggested that the neural representation of hand actions is organized based on the extent of sociality and transitivity of these actions. Recently, in a study with a large set of natural images of actions, Tucciarelli et al. (2019) compared the similarity in neural representation with their semantic similarity obtained from behavioral ratings. They showed that the neural organization of observed actions in the lateral occipito-temporal cortex is correlated with the behavioral similarity judgments.

Investigations to the representational content of parietal action selective regions are fewer in number, and their results are more subject to debate. Shmuelof and Zohary (2006, 2008) have proposed an effector dependent representation of hand actions in the anterior intraparietal cortex, while others have argued actions are represented in an effector independent manner in the inferior parietal lobe (Jastorff et al., 2010) as well as superior parietal lobe and intraparietal sulcus (Vingehoets et al., 2012). One study (Jastorff et al., 2010) suggested that action representations in the parietal cortex are related to the direction of the action relative to the body.

Looking at the entire visual system, Tarhan and Konkle (2020) used an encoding model to predict responses to natural videos. The feature space of their model captured the body parts involved in an action and the action target. Based on the voxel tunning, they suggested that five large-scale networks exist in the human brain that represent actions based on their sociality and the spatial extent of their interaction envelope. All these studies have employed natural videos of actions in which often a whole scene and objects are present. Employing natural stimuli may lead to many confounding factors and difficulties in interpreting the results. Even though some of these studies have taken steps to make sure low-level features are not contributing to their results (Tucciarelli et al., 2019; Tarhan and Konkle, 2020), there is a possibility their results may be related to the presence of special types of objects, scene contexts and semantic relations between them (Kourtzi and Kanwisher, 2000; Senior et al., 2000; Johnson-Frey, 2004; Buxbaum et al., 2006; Wurm et al., 2012; Schubotz et al., 2014; el-sourani et al., 2017; Wurm and Schubotz, 2017; Wurm et al., 2017a; Leshinskaya et al., 2018).

Here, employing fMRI multivoxel pattern analysis, we decoded the position invariant representation of human actions in point-light display (PLD) format. We used controlled stimuli in PLD format to restrict the visual information to the bodily movements and identify regions supporting abstract action representations. Using these controlled action stimuli, we localized regions where action decoding was robust to changes in position. Employing PLD stimuli also allowed us to determine the position and movements of individual limbs, to devise objective measures for constructing similarity between actions. Using these objective measures of similarity as well as subjective measures derived from behavioral experiments, we characterized the representational content of action selective regions.

## Materials and Methods

### Participants

Twenty-five subjects (14 females, 20-38 years of age) took part in the fMRI experiment and 15 subjects (13 females, 25-38 years of age) took part in the behavioral experiment. All subjects were healthy and right-handed with normal visual acuity. They gave written informed consent and received payment for their participation. The experiments were approved by the ethics committee on the use of human subjects at the Iran University of Medical Sciences. Three fMRI subjects were later excluded due to excessive motion (see data analysis for more details) and the final analyses included the remaining 22 subjects.

### Functional MRI experiment

In this experiment, we used videos of 10 different human actions in Point Light Display (PLD) format: Crawl, Cycle, Jumping Jack (henceforth referred to as Jump), Peddle, Play tennis, Salute, Spade, Stir, Walk, and Wave (Figure 1). These actions were selected from a larger set of human action videos in PLD format provided by Vanrie and Verfaillie (2004).

**Figure 1.**
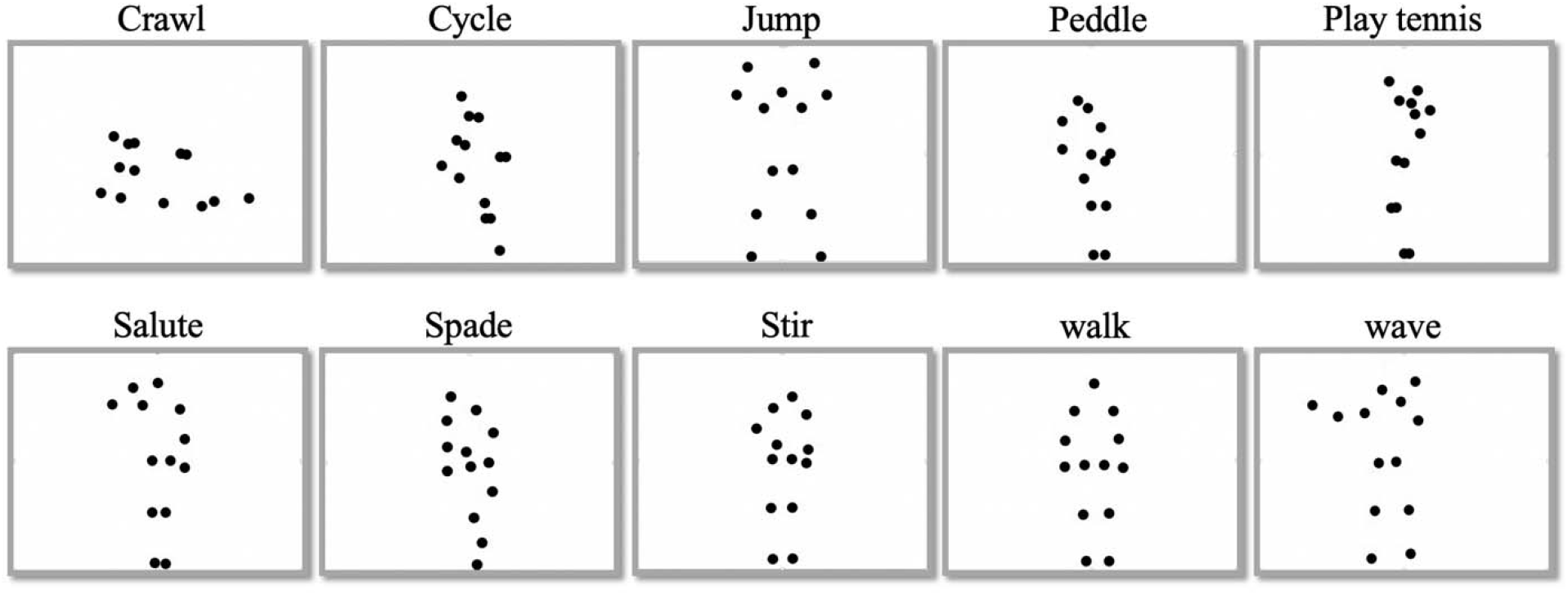
Representative frames of the 10 human action videos in PLD format used in the experiment.

We used a block design paradigm. Each run included twenty blocks with two blocks for each human action video. In one of the blocks for each action, the video was presented in the upper visual hemifield, and in the other, it was presented in the lower visual hemifield. The presentation order of the stimuli was counterbalanced across runs. Each block lasted 8 seconds. There was an 8-second blank period in the beginning, end, and between stimulus blocks of each run. Each run lasted 328 seconds. Each subject completed one session of 10 runs.

In each stimulus block, there were four repetitions of the same human action video. Each repetition lasted 1.5 seconds (frame rate 30), followed by a 0.5 second blank period. In a randomly chosen repetition in each block, the size of the dots became 50% larger for one second. The point-lights, subtending ∼0.25 degrees of visual angle, were in white color against a black background. The center of the stimuli was presented ∼4 degrees of visual angle above/below the fixation point. The size of the area that the dots occupied during movements was between 0.94 and 5.62 degrees in width and between 2.38 and 7.7 degrees in height.

The fixation point was a red circle of size ∼0.25 degrees of visual angle. Subjects were instructed to fixate on the fixation point at the center of the screen, watch the videos, detect the change in the size of the point lights, and report it by pressing a response key with their right index finger.

### Behavioral experiment

To capture the similarity between our human action stimuli, we conducted a behavioral experiment based on an inverse multi-dimensional scaling (IMDS) method proposed by (Kriegeskorte & Mur, 2012). At the beginning of the experiment, one snapshot from each of the ten action videos was presented along with a number 1-10 in two vertical columns at the left border of the screen. On a separate screen, they were provided with numbered action videos, and they could watch them as many times as they needed. Participants were asked to rearrange the snapshots on the surface of a gray circle (drag and drop using the mouse) according to the perceived similarity of their corresponding videos. The arrangement was performed based on subjective judgments of overall similarity, and no other instructions were given regarding the specific aspects they needed to focus on for the arrangement. A behavioral dissimilarity matrix was obtained based on the final Euclidean distances on the circle for each participant.

### MRI methods

MRI data were collected at two centers: the school of Cognitive Sciences, Institute for Research in Fundamental Sciences (Tehran, Iran) using a Siemens 3T Tim Trio MRI scanner and a 32-channel head coil and the National Brain Mapping Lab (Tehran, Iran) using a Siemens 3T Prisma MRI scanner and 20-channel or 64-channel head coil. We were forced to switch MRI equipment mid-experiment due to scanner/coil malfunction (data of 5 subjects were recorded at the school of Cognitive Sciences, and data of 20 subjects were recorded at National Brain Mapping Lab). We moved scanners to ensure that high-quality data was collected for all subjects. Similar protocols were used across scanners, and the data recorded with different equipment were comparable.

Subjects viewed the visual stimuli through a back-projection screen, and the task was presented using MATLAB and Psychtoolbox-3 (Brainard, 1997). Functional images were obtained using a T2^*^-weighted single-shot gradient-echo EPI sequence with a repetition time (TR) of 2 s, echo time (TE) 26, 90° flip angle, 30 transverse slices, and a voxel size of 3 × 3 × 4 mm^3^. A high-resolution T1-weighted structural scan was also acquired from each participant using an MPRAGE pulse sequence (TR = 1800 ms, TE = 3.44 ms, inversion time = 1100 ms, 7° flip angle, 176 sagittal slices, and 1 × 1 × 1 mm^3^ isotropic voxels).

### Data analysis

Freesurfer (https://surfer.nmr.mgh.harvard.edu) and FS-FAST (Dale et al., 1999) were employed for data preprocessing and general linear model (GLM) analysis. Preprocessing of fMRI data included motion correction, slice timing correction, linear and quadratic trend removal, and no spatial smoothing. Subjects with excessive head motion (more than 1.5 mm within a run and 4 mm across runs) were excluded (3 subjects). Functional data were then resampled to the cortical surface of individual subjects.

A run-wise GLM analysis was performed to obtain the beta values and their corresponding t statistic for each human action and each presentation hemifield in each vertex (10 human actions in the upper visual field and 10 human actions in the lower visual field leading to 20 regressors). Linear and quadratic polynomial nuisance regressors and external regressors from the estimated head movements were also included.

Wang atlas of visual topography (Wang et al., 2014) was used to localize retinotopic areas for each subject: V1, V2, V3, hV4, VO1, VO2, MST, hMT, LO2, LO1, V3a, V3b, IPS0, IPS1, IPS2, IPS3, IPS4, IPS5, SPL1 (Figure 3).

### Multivariate pattern analysis (MVPA)

In-house MATLAB codes, LibSVM (Chang and Lin, 2011), and CoSMoMVPA (Oosterhof et al., 2016) toolboxes were employed for searchlight and ROI analyses.

#### Searchlight analysis

We performed action classification using a surface-based searchlight procedure to obtain a map of classification accuracy (Oosterhof et al., 2011). In a leave-one-run-out cross-validation procedure, samples (t-statistics of vertices) were partitioned to train and test sets. A linear Support Vector Machine (SVM) was employed to perform pairwise brain decoding classification for each presentation position (upper/lower visual hemifield) in individual brains with a searchlight circle of 100 vertices on the surface. The decoding accuracy of a searchlight was calculated as the average of all within position pairwise classification accuracies. The resulting maps were then resampled to a common surface for the group-level statistical analysis.

To determine regions showing position tolerance, we performed cross-position decoding by training a classifier to discriminate pairs of actions presented in one position and testing its accuracy in classifying the same pair of actions in the other position. A cross-position decoding accuracy greater than the chance level would indicate position tolerance.

To obtain group-level statistics, p-values of classification accuracies were computed using a binomial test, corrected for false discovery rate (Benjamini and Hochberg, 1995) at level q1. The second-level p-value for each vertex was then determined as

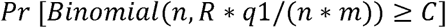

With *C* denoting the number of participants for which that vertex was significant, *R* denoting the sum of the counts of C across all vertices, *n* denoting the number of participants, and *m* denoting the number of vertices. Under the null hypothesis, C has a binomial distribution with size n and a probability that is approximately bounded by *R* * *q*1/(*n* * *m*). Finally, the derived second-level p-values were thresholded at the FDR level q2 to acquire the significant vertices at the group-level (McMahon et al., 2019). This approach is more appropriate than the t-test for comparing classification accuracies against chance level (See Allefeld et al., 2016 for a discussion of why the t-test is not suitable in such cases).

From the searchlight group-level statistical analysis, we obtained clusters with significant cross-positions classification accuracy. Contiguous clusters that passed the threshold were selected as ROIs in which decoding of actions could be generalized across positions and used in the next analyses to explore their representational structure. To ascertain that the ROI selection and further analyses are statistically independent and prevent double-dipping, we used a leave-one-subject-out approach (Etzel et al., 2013). The group-level statistical analysis was applied to the searchlight results of all subjects except one, and the resulting ROI was projected on the left-out subject to select the ROI in that subject. The procedure was repeated for each subject. The same clusters were found consistently across all iterations.

#### Comparing within and cross-position classification accuracies

To directly compare within and cross-position classification accuracies, we performed SVM classifications within each ROI obtained from the leave-one-subject out analysis described above. The procedure for obtaining within- and cross-position classifications was the same as the searchlight analysis with the addition of feature selection. Different ROIs don’t include the same number of vertices. This variation in the number of vertices across ROIs could influence classification accuracies. To avoid this potential confounding factor, the pairwise classifications were performed using the 100 most informative vertices in each ROI. To select the most informative vertices, a t-test was applied on the training set, and 100 vertices with the lowest p-values for discriminating between the conditions of interest in the training set were chosen (Mitchell et al., 2004). It is noteworthy that the classification results didn’t qualitatively differ without feature selection.

#### Exploring the representational content of the ROI

To explore the representational content of the ROIs, we constructed models based on the body-part movements and behavioral ratings and examined the correlation between these models and the cross-position decoding accuracies for each ROI.

To obtain the body-part movement model, we divideded the point lights into five groups: trunk (5 dots), left hand (2 dots), right hand (2 dots), left leg (2 dots), and right leg (2 dots). We obtained the sum of the displacements of point lights in each group for each action video. From the total displacements in each group, we obtained a five-element vector. Then, the Euclidean distance between these five-element vectors was used to obtain a dissimilarity matrix based on the body-part movement. This body-part movement model includes information on both the pattern of body-part movement and the average of their movement for each action. Hence, for further investigation, we examined two other models: body-movement-pattern and body-movement-average. To obtain the body-movement-pattern model for each action, we demeaned the corresponding five-element vector of the body-part movement model and obtained the Euclidean distance between these new demeaned vectors to build a dissimilarity matrix. The difference between the average values of body-part movement vectors was also used to obtain the body-movement-pattern model.

To obtain the behavioral dissimilarity model, the results from the behavioral experiment were used. In the behavioral experiment, the distance between stimuli indicates their (dis)similarity. The obtained dissimilarity matrix for each subject was normalized by dividing each value in the matrix by the maximum value. The pooled behavioral dissimilarity matrix was then computed as an average of individual normalized behavioral dissimilarity matrices.

To obtain the correlation between the models and the action representations in each ROI, the off-diagonal of the matrix obtained from the pairwise cross-position decoding accuracies was vectorized. The off-diagonal of the dissimilarity matrices from the models were also vectorized. The Kendall rank correlation between dissimilarity vectors and decoding accuracy vectors was then calculated, and the correlation values were compared to characterize the representational content of each ROI.

#### ROI similarity analysis

We also performed the cross-position decoding accuracies for Wang’s ROIs using a similar procedure as that used for the ROIs obtained from the searchlight analysis. Following a leave-one-run-out approach, a classifier was trained to decode pair of actions presented in one position and tested with the same pair of actions in another position. We applied a t-test on the training set to choose 100 vertices with the lowest p-values for discriminating between the actions of interest in each ROI. Then, to investigate the representational similarity between ROIs (Wang’s ROIs and ROIs obtained from searchlight analysis), correlations were computed between the vectorized matrices of pairwise cross-position decoding accuracy. One minus these correlation values were used to obtain the distance between the ROI pairs, and these distances were used to construct an ROI dissimilarity matrix. The ROI dissimilarity matrix was first computed for individual subjects and then averaged across subjects to acquire group level ROI dissimilarity matrix (Figure 7A).

Split-half reliability of ROI dissimilarity matrix was also evaluated; we randomly divided subjects into two equal groups and correlated corresponding group-level ROI dissimilarity matrices as a measure of reliability. This measure was calculated for 10000 random split-half divisions and averaged to produce the final reliability measure. To examine the significance of this reliability measure, we obtained the bootstrapped null distribution of reliability by random shuffling of the labels in the correlation matrix separately for the two split-half groups and calculating the reliability for 10000 random samples.

An MDS analysis was then performed on this ROI-dissimilarity matrix, and the first two dimensions that captured most of the variance were used to produce an MDS diagram in which the distance between each pair of ROIs represents the similarity between them (Vaziri-Pashkam & Xu, 2018). The squared correlation (*r*^*2*^) between two-dimensional distances in the MDS plots and the original distances in the multi-dimensional space was used to quantify the variance explained by the first two dimensions. We then fit two regression lines employing the total least square method, with one line passing through the positions of the occipito-temporal regions and the other passing through the positions of parietal regions. The variance explained by the fitted lines was also measured; two-dimensional distances between regions based on the predicted positions on the lines were obtained, and the *r*^*2*^ between these distances and the original distances based on the input matrix to the MDS analysis was calculated. Similarly, we computed the *r*^*2*^ between distances based on the MDS plots and line predictions to determine the variance explained by the two lines on the MDS plot.

## Results

In the present study, we aimed to identify regions containing position tolerant representations of actions and investigate the representational content of these regions using fMRI multi-voxel pattern analysis. A stimulus set of ten different human action videos in PLD format (Figure 1) was presented at two different positions in the visual display (upper and lower visual hemifields), and t-values were extracted in each voxel of the brain for each of the actions and each position. Employing a support vector machine classifier, within- and cross-position decoding was then applied following a searchlight approach across the whole brain.

### Action decoding across the whole brain

To identify regions that showed selectivity for our action stimuli within each presentation position (upper/lower visual fields), classifiers were trained and tested with fMRI responses for stimuli presented within the same position. To identify regions showing position tolerance, we performed a cross-position classification. Classifiers were trained with fMRI responses to stimuli presented at the upper visual field and tested with fMRI responses to stimuli presented at the lower visual field and vice versa. Both within- and cross-position classification were applied using a surface-based searchlight method.

Figure 2A and B depict the result of the group analysis (q1 = 0.05 and q2 = 0.01) for within- and cross-position classification accuracy, respectively. Comparison of these maps reveals that a subset of the regions that show above chance within-position classification also demonstrate generalization across positions. The group analysis showed significantly above chance cross-position classification accuracy in the left and right lateral occipito-temporal cortex, right intraparietal sulcus, and right post-central gyrus (Figure 2B).

**Figure 2.**
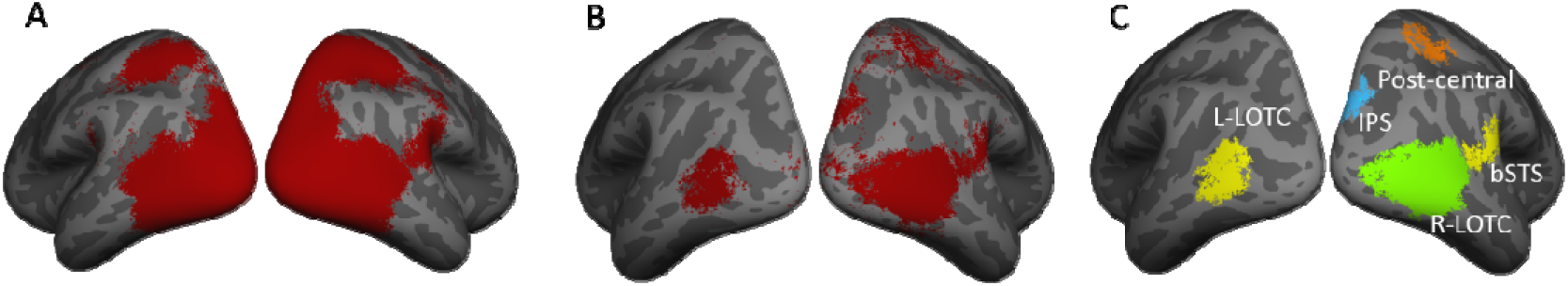
Results of searchlight classification accuracy ***A***, Significance maps of group analysis for searchlight within-position classification accuracy (thresholded at *q1* = 0.05 and *q2* = 0.01) ***B***, Significance maps of group analysis for searchlight cross-position classification accuracy (group thresholded at *q1* = 0.05 and *q2* = 0.01) ***C***, five Clusters with significant cross-position classification accuracy used as ROIs (note that the exact ROIs differ slightly across iterations of the leave-one-subject-out procedure).

**Figure 3.**
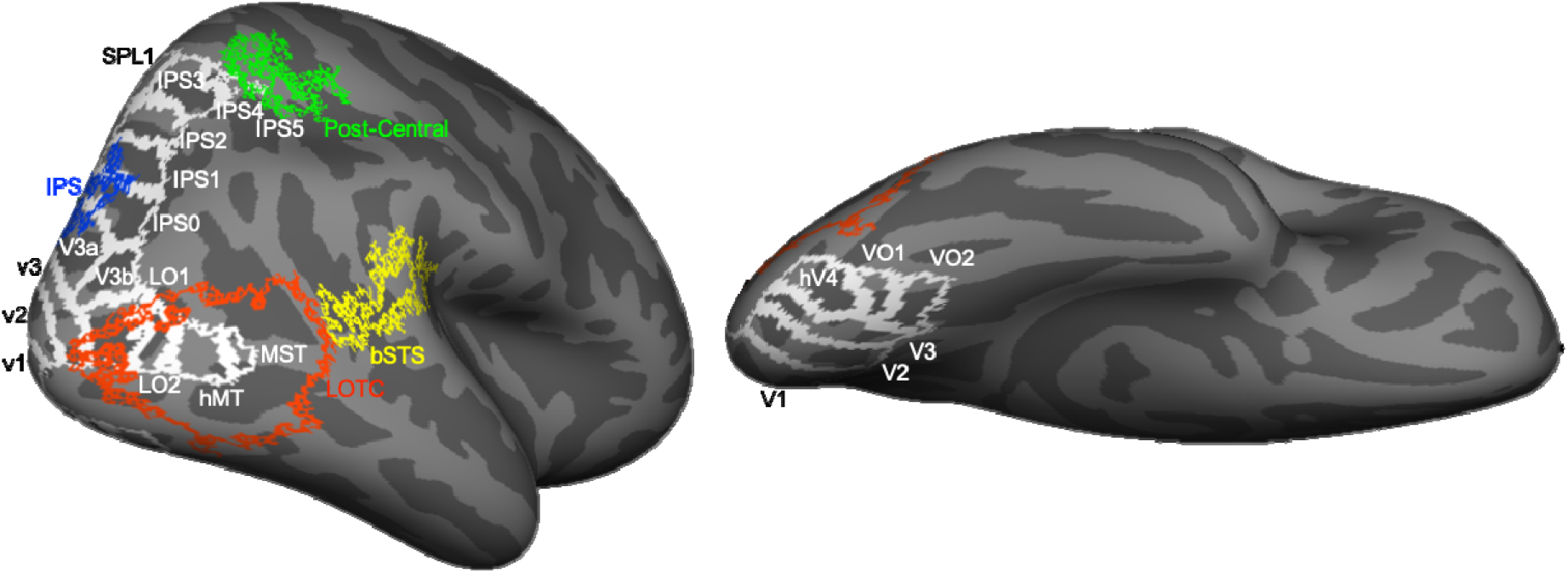
Inflated brain surface from a representative participant showing the ROIs examined in lateral and ventral views with white outlines for Wang’s ROIs and colorful outlines for functional ROIs obtained from searchlight analysis.

### Action decoding in ROIs

To investigate the extent to which changes in position affect the representations in individual regions, we used a leave-one subject out procedure (see methods) to extract ROIs with above chance cross-decoding accuracy. Five clusters (Figure 2C) were selected based on a searchlight analysis on all but one subject, and the selected clusters were used as ROIs for the left-out subject.

A linear SVM was employed for within- and cross-position classification within each ROI, using the 100 most informative vertices in each ROI (Mitchell et al., 2004). Figure 4A illustrates the results of within- and cross-position decoding accuracy for each of the ROIs. All ROIs had a significant decoding accuracy for cross- and within-position (group analysis with (q1 = 0.01, q2= 0.01). In the left and right lateral occipito-temporal and parietal ROIs, cross-position decoding accuracy was significantly lower than within-position decoding accuracy (paired t-test, p < 0.0075, FDR corrected q = 0.01), while this difference was not significant in bSTS and Post-Central regions (paired t-test, p > 0.13, FDR corrected q = 0.01). These results suggest a reduction in position sensitivity from posterior to anterior regions in both occipito-temporal and parietal cortices. To visualize this finding on the whole brain, we divided the cross-position by within-position classification accuracy in all vertices with above chance within-position accuracy. The resulting map averaged across subjects is depicted in figure 4B. In line with the ROI analysis, this map demonstrates an increase in position tolerance from posterior to anterior regions in both occipito-temporal and parietal cortices.

**Figure 4.**
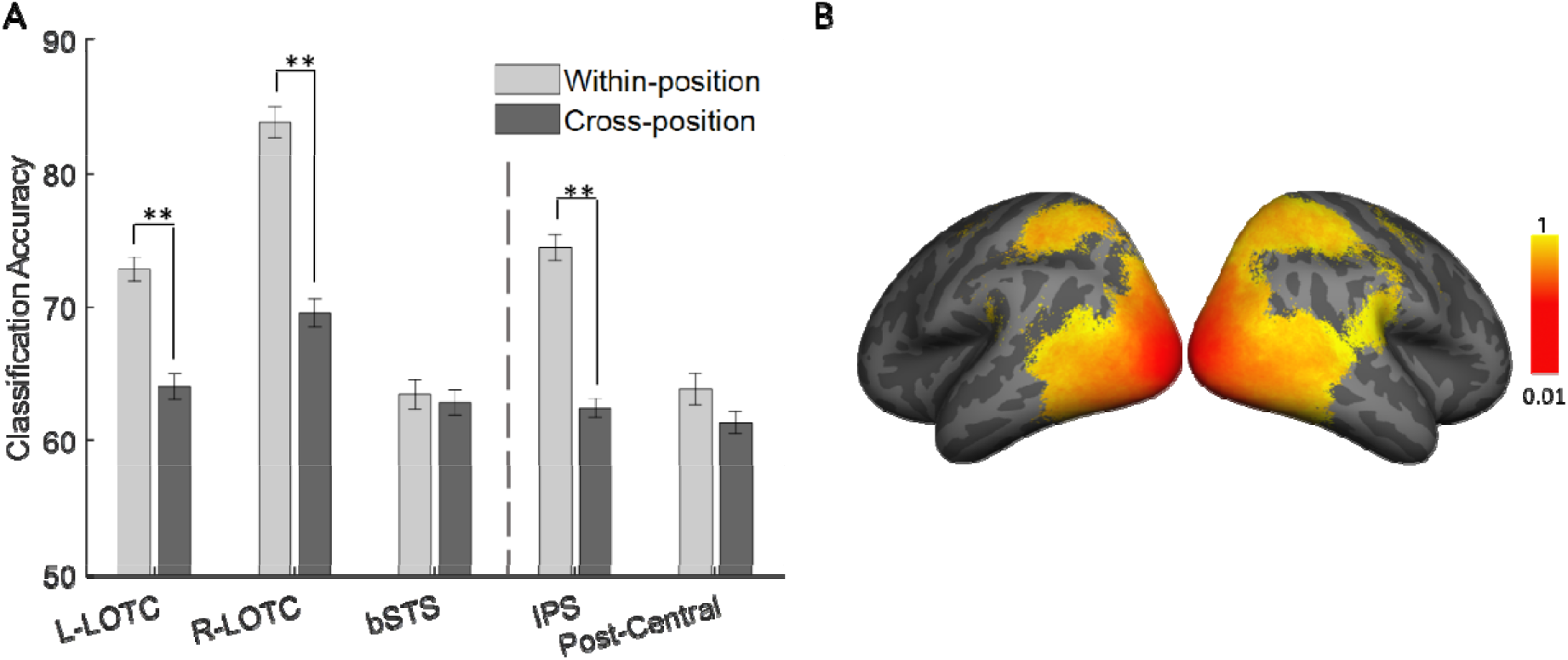
***A***, Within and cross-position decoding accuracy for the ROIs obtained from searchlight analysis. Error bars indicate standard errors of the means (** paired t-test, p < 0.01, FDR corrected). The vertical dashed line separates occipito-temporal ROIs from parietal ROIs ***B***, Maps of cross-position divided by within-position classification accuracy obtained from searchlight analysis in vertices with significant within-position classification accuracy.

#### Revealing the Representational content of the ROIs

To uncover the principles that govern the organization of action representations in each ROI, we used a representational similarity analysis (RSA, Kriegeskorte et al., 2008). We constructed models based on the body-part movements and behavioral similarity ratings performed on the videos by human observers (see methods). We then used correlation analysis to investigate the similarity of action representations between models and ROIs according to their cross-position pairwise decoding accuracies. The reliability of the behavioral dissimilarity matrices was high (Cronbach Alpha = 0.7654). Models of stimuli (distance matrices obtained from models) were also compared with each other; the Kendall correlation was calculated, and its significance was examined applying permutation tests. The behavioral and body-part models were not significantly correlated (Kendall correlation = 0.0394, p-value = 0.2562).

Figure 5 shows the Kendall correlations between the models and cross-position pairwise decoding accuracies across participants for body-part movement and behavioral rating models.

**Figure 5.**
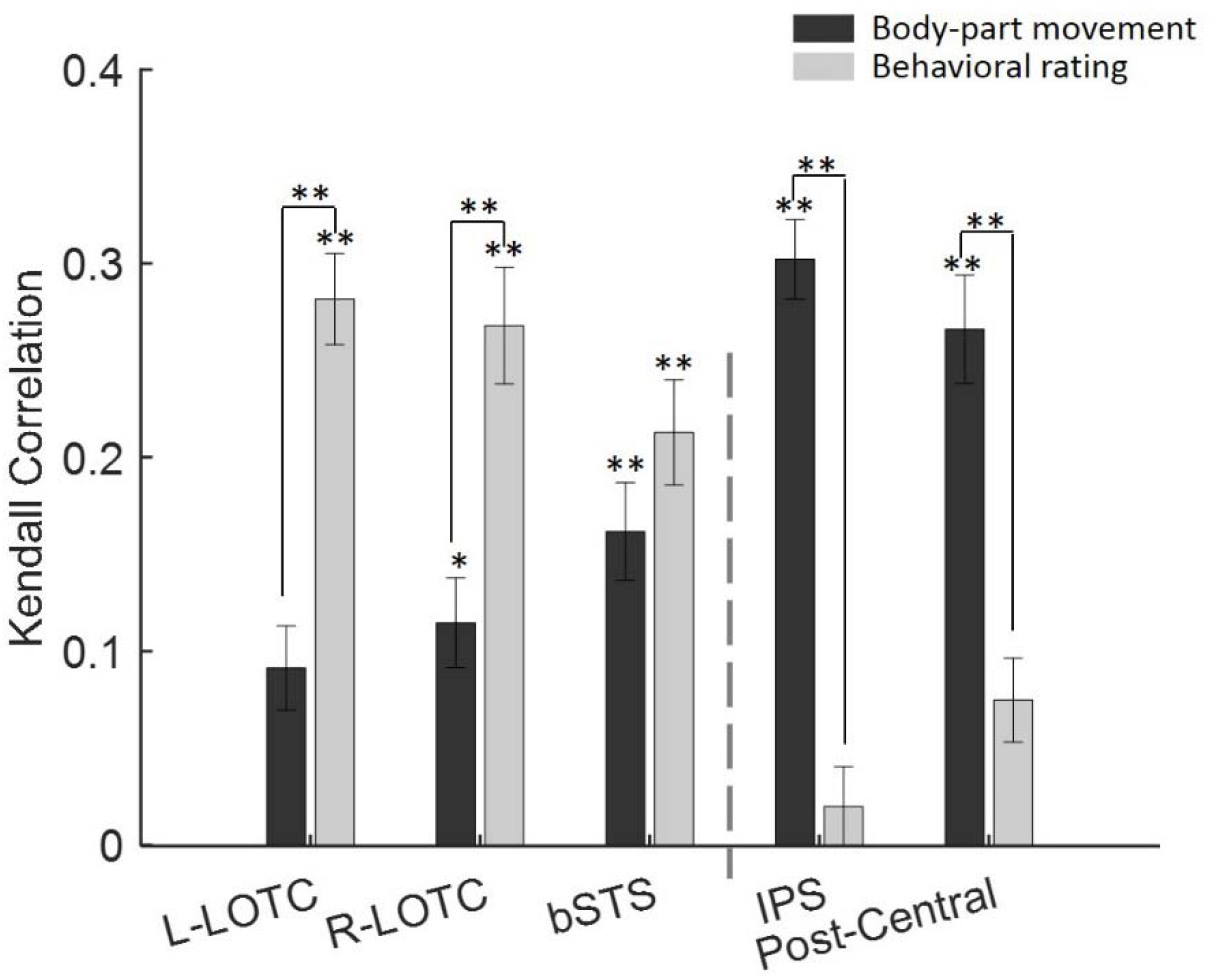
Kendall correlation body-part movement (dark gray) and behavioral rating models (light gray) and cross-position pairwise decoding accuracies for the individual ROIs. Error bars indicate standard errors of the mean (permutation test or paired t-test, \**p* < 0.05, \*\**p* < 0.01, FDR corrected). The vertical dashed line separates occipito-temporal ROIs from parietal ROIs.

Significant correlations were found with the body-part model in all ROIs except for L-LOTC (permutation test, *p* < 0.05 for R-LOTC and *p* < 0.01 for bSTS, IPS, Post-Central) and with the behavioral rating model in all ROIs except for IPS and Post-Central (permutation test, *p* < 0.01).

A two-way repeated-measures ANOVA with model (body-part movement and behavioral rating) and ROI as independent factors and correlation coefficient as the dependent variable was performed. The effect of ROI (*F*(4,84) = 0.9303, *p* = 0.4504), and the effect of model (*F*(1,84) = 0.7636, *p* = 0.3921) were not significant while the interaction between the two (*F*(4,84) = 36.8, *p* < 0.0001) was significant. Individual comparisons within ROIs showed that correlation coefficients were greater for the behavioral rating model in L-LOTC and R-LOTC and for the body-part movement model in IPS and post-central (all *t*(21) > 3.7117, all *p* < 0.0016, FDR-corrected).

We also constructed models based on the average of movement across all body-parts (body-movement-average) and based on the pattern of movements in body parts (body-movement-pattern) separately (see methods). Figure 6 shows the average Kendall correlation between the models and cross-position pairwise decoding accuracies across participants for body-movement-pattern and body-movement-average models. As figure 6 shows, all ROIs showed correlations with the body-movement-pattern model (permutation test, *p* < 0.05 for L-LOTC and R-LOTC, *p* < 0.01 for bSTS, IPS and Post-Central, FDR corrected, 5 comparisons), but body-movement-average models only showed correlations with the dorsal ROIs (permutation test, *p* < 0.01, FDR corrected) and showed no significant correlation with the lateral ROIs (permutation test, p > 0.05, FDR corrected).

**Figure 6.**
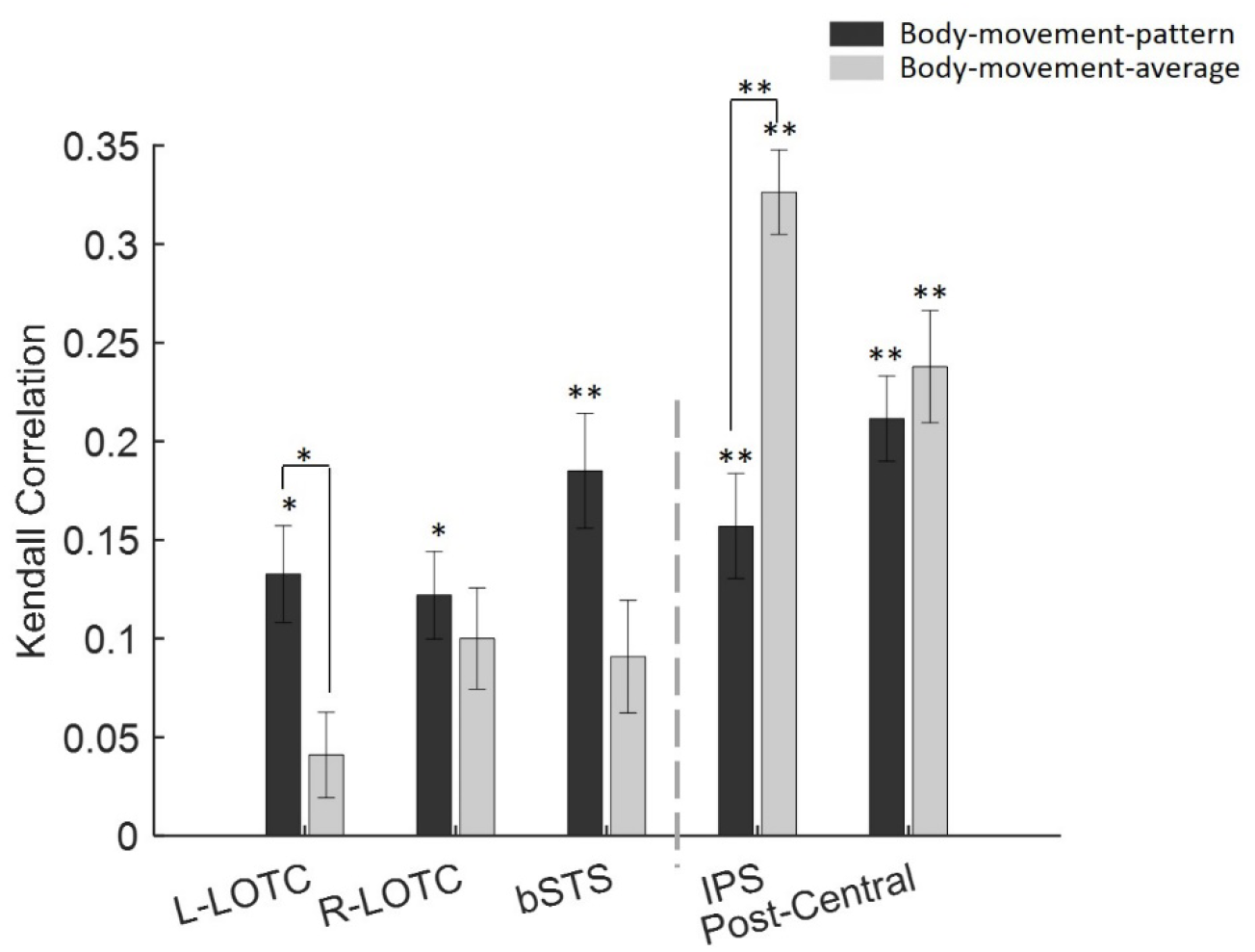
Kendall correlation between body-movement-average (light gray) and body-movement-pattern (dark gray) models and cross-position pairwise decoding accuracies for individual ROIs. Error bars indicate standard errors of the mean (permutation test, \**p* < 0.05, \*\**p* < 0.01, FDR corrected). The vertical dashed line separates occipito-temporal ROIs from parietal ROIs.

**Figure 7.**
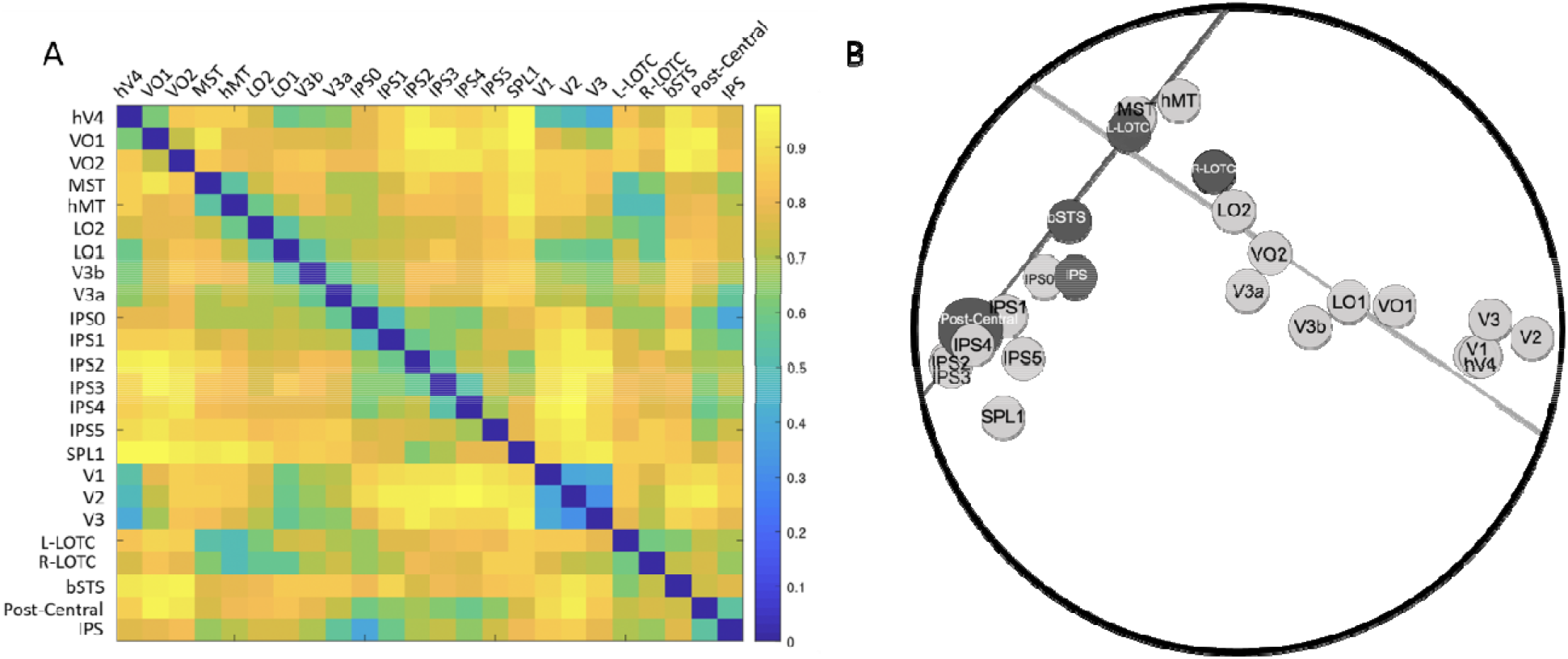
***A***, ROI dissimilarity matrix based on the correlation distance between their cross-position decoding accuracy for Wang’s ROI and ROIs obtained from searchlight analysis. Lighter colors show higher correlations ***B***, MDS plot of Wang’s ROIs (light gray) and ROIs obtained from searchlight analysis (dark gray) and a total least square regression line through the occipito-temporal regions (the light gray line) and another through the parietal regions (the dark gray line). The MDS plot and the lines could capture 0.5522 and 0.4798 percent of the variance of the ROI dissimilarity matrix.

A two-way repeated measure ANOVA with model and ROI as independent factors and correlation coefficient as the dependent variable showed no significant effect of model (*F*(1,84) = 0.0117, *p* = 0.9148) while the effect of ROI (*F*(4,84) = 22.5280, *p* < 0.0001), and the interaction between the two (*F*(4,84) = 11.5105, *p* < 0.0001) were significant. The significant effect of ROI points to greater correlation coefficients in dorsal ROIs. Individual comparisons within ROIs showed that correlation coefficients were greater for the pattern model in L-LOTC (*t*(21) = 2.6553, *p* = 0.0370, FDR-corrected) and for the average model in IPS (*t*(21) = 4.9394, *p* = 3.4583e-04, FDR-corrected).

We also compared the behavioral rating model with the average and pattern models applying two separate two-way repeated measure ANOVAs with model and ROI as independent variables and correlation coefficient as the dependent variable.

Its result for the body-part-average and behavioral rating models showed no significant effect of ROI (*F*(4,84) = 1.1066, *p* = 0.3589), and model (*F*(1,84) = 0.3031, *p* = 0.5878) while the interaction between the two was significant (*F*(4,84) = 36..4416, *p* < 0.0001). Individual comparisons within ROIs showed that correlation coefficients were greater for the behavioral rating model in L-LOTC and R-LOTC and for the body-movement-average model in IPS and Post-Central (all *t*(21) > 3.5686, all *p* < 0.0018, FDR-corrected).

For the body-part-pattern and behavioral rating models results showed no significant effect of model (*F*(1,84) = 0.4335, *p* = 0.5174) while the effect of ROI (*F*(4,84) = 8.8253, *p* = 0.0001), and interaction between the two (*F*(4,84) = 37.6654, *p* < 0.0001) were significant. Individual comparisons within ROIs showed that correlation coefficients were greater for the behavioral rating model in L-LOTC and R-LOTC and for the body-movement-pattern model in IPS and Post-Central (all *t*(21) > 4.8254, all *p* < 0.0001, FDR-corrected).

In sum, these results reveal dissociations between the classification accuracies in the lateral and dorsal ROIs. In lateral ROIs, the cross-position decoding accuracies are more correlated with behavioral ratings than any of the body-part movement models, while those in dorsal ROIs are more correlated with the objective movement models.

#### ROI similarity analysis

In addition to the 5 clusters obtained from the searchlight analysis, we obtained the cross-position decoding accuracy of retinotopic regions of interest across occipito-temporal and parietal cortex (see methods). We calculated cross-position classification accuracies for these ROIs following a similar procedure as that used for the ROIs obtained from searchlight analysis. Then, to examine the representational similarity across all ROIs (including retinotopic ROIs and ROIs obtained from searchlight analysis), we computed the one minus the correlation between their cross-position decoding accuracy (Figure 7A). The resulting ROI dissimilarity matrix had high split-half reliability of 0.8346, significantly higher than chance (bootstrapped null distribution 95% CI [−0.0011,0.0012]). To visualize region-wise distances, we then applied an MDS analysis on this matrix. Figure 7B depicts the first two dimensions that captured most of the variance (*r*^*2*^ = 0.5522). These results are consistent with the representational similarity analysis and further suggest the presence of two separate networks for processing actions. One starting from the early visual cortex going to the lateral surface of the brain, and another splitting from this pathway towards the parietal cortex with the bSTS region landing right between the two pathways. This observation was quantified through fitting two total least square lines to the points on the MDS plot: one line was passed through occipito-temporal regions (V1, V2, V3, hV4, VO1, VO2, MST, hMT, LO2, LO1, V3b, and V3a from the wang atlas, and L-LOTC, R-LOTC, and bSTS from our ROI analysis) and the other line was passed through parietal regions (IPS0, IPS1, IPS2, IPS3, IPS4, IPS5, and SPL1 from wang atlas and IPS and Post-Central from our ROI analysis). These lines could explain 0.7950 percent of the variance of the positions of the regions on the MDS plot and 0.4798 percent of the variance of the full multi-dimensional space.

## Discussion

In this article, we used point-light displays of human actions presented in either the upper or the lower visual fields and examined the extent of position tolerance as well as the representational content of action selective regions in the human brain. Using cross-position decoding analysis, we found that regions in the Left and right lateral occipitotemporal cortex, right intraparietal sulcus, and right post-central gyrus contain position tolerant representations of action stimuli. Additionally, we investigated the representational content of these regions and found that representations in parietal regions were related to the movements of the body-parts while those in occipitotemporal regions were more related to human subjective judgments about actions.

Our results, for the first time, provide a comprehensive understanding of position-tolerant action representations in the human visual cortex. Our findings in the lateral occipital cortex are consistent with those of Grossman et al. (2010) that applied fMRI adaptation and showed position tolerant representation of actions in STS. However, they failed to find other position invariant regions found in our study, especially the regions in the parietal cortex. This discrepancy may be related to differences in design. Our block design paradigm and the use of SVM classifiers may have provided us with a higher power and sensitivity to detect regions that contain position-tolerant action representation (Coutanche et al., 2016). Roth and Zohary (2015) investigated position invariance during observation of natural tool-grasping clips using MVPA (correlation and cross-decoding classifier). They observed a gradual loss of position information accompanied by enhancement of hand/tool identity along the posterior-anterior axis in the dorsal stream. This result is consistent with our finding for the parietal cortex, but they didn’t observe such a gradient in the lateral stream. Their focus on grasping actions may have lowered their chance of observing the full extent of position tolerance in the lateral occipito-temporal cortex.

Most regions that contained position tolerant representations of actions in our study were located in the right hemisphere. This laterality toward the right hemisphere is in line with the results of previous studies. Beauchamp et al. (2003) investigated human motion versus tool motion (natural and PLD video clips). They found that category-related responses were lateralized with a significantly greater number of tool-preferring voxels in the left hemisphere and a significantly greater number of human-preferring voxels in the right hemisphere. Grosbras et al. (2012) performed a meta-analysis on three categories of human motion including the movement of the whole body, hands and face. The conjunction of the three meta-analysis maps showed convergence in the right posterior superior temporal sulcus and the bilateral junction between middle temporal and lateral occipital gyri. Right laterality in parietal regions has also been previously reported for observed limb movements (Grosbras et al., 2012; Pelphrey et al., 2005).

After identifying the position-tolerant regions, we investigated their representational content. In occipito-temporal cortex, we found that a model based on the behavioral ratings of similarity was the best predictor of the cross-position pattern classification accuracies. These results are consistent with a study by Tucciarelli et al. (2019) that used natural images of different actions and showed that responses in the LOTC were correlated with the behavioral ratings of semantic similarity between actions. Similar to our results, they also showed a lower correlation in LOTC with a model based on body movements. On the other hand, they failed to find a significant correlation with a body part model in the parietal cortex. This could be because they used human ratings of body part movements as opposed to the objective measures we used based on actual measures of body position. Humans may fail to consider the movement of all body parts and only focus on just the main effector of action for their ratings. In addition, they also employed an event-related design that has less power than our block design paradigm. Thus, our results extend their findings and further elucidate the differences between parietal and occipito-temporal action representations.

In a study with a large set of human action videos, Tarhan and Konkle (2020) used behavioral rating to construct an encoding model to investigate action representations in the human brain. Their behavioral ratings included questions related to body parts involved in the action as well as the action target. Using a clustering analysis, they divided voxels into four network, suggesting that one is driven by the social aspects of actions and four are driven by the scale of the interaction envelope. Consistent with our study, their clusters spanned both occipitotemporal and parietal regions, with notable differences in response profile between the two networks. However, it is difficult to be more specific in comparing their results to our findings due to methodological differences. They used natural videos of everyday scenes containing actions as stimuli. Other than human bodies or body parts performing the action, their stimuli also contained objects and other scene elements. The use of natural stimuli makes it difficult to determine if the clusters reflect responses to actions or the scene elements correlated with the actions. Also, since we used a block design paradigm and aimed at determining the extent of position invariance in action selective regions, we could only collect data from a small set of actions. Future studies with a larger set of controlled PLD stimuli could test the extent to which sociality and interaction envelope determine the responses in position invariant action selective regions in the brain.

Investigating the representations of actions in the dorsal stream, Buccino et al. (2001) suggested that dorsal regions represent actions based on action effectors. Jastorff et al. (2010), Abdollahi et al. (2013), and Vannuscorps et al. (2018), on the other hand, provided evidence that action representations in dorsal regions are organized based on the category of actions independent of the effectors. Here, we show that a model based on body-part movement shows high correlations with the pattern of classification accuracies in the dorsal stream regions. Our body-part movement model was constructed based on the movement of all body parts, including the trunk and the limbs. This model included more information than just the action effector. High correlations of this model with the classification accuracies in the dorsal regions suggests that dorsal regions may contain a more holistic representation of body parts during action observation.

Even though lateral regions show higher correlations with the subjective model than the body part model, their correlation with the body part movement was still above zero. Therefore lateral regions show signatures of both objective and subjective measures. Nevertheless, more detailed analysis of the body part model separating the patterns of movement from average movements revealed further differences between the dorsal and lateral regions. While the dorsal regions showed sensitivity to both pattern and average of movements, the lateral regions showed only a correlation with a model based on patterns of movement. With the limited number of stimuli in our set, we cannot perform a more nuanced analysis of the differences in the sensitivity to various body parts between dorsal and lateral regions. Future experiments with an expanded set of PLD stimuli could reveal potential differences between the two networks in their body part representations.

In the domain of object representations, recent studies have revealed that both dorsal and ventral stream regions contain object information independent of the position of objects (Vaziri-Pashkam and Xu, 2019; Almeida et al., 2018), but data-driven analysis of object representation still reveals the separation of these dorsal and ventral regions in their representational (Vaziri-Pashkam and Xu, 2019). Here, focusing on action observation, we showed that action representations become progressively less position-dependent from posterior to anterior along both dorsal and lateral pathways. Moreover, employing either data-driven (looking at ROI similarities) or model-based approaches, we observed a clear distinction between representational content in two pathways. Looking more closely at the arrangement of ROIs in the MDS plot, we can see that ROIs along the lateral surface follow a continuous trajectory from early visual areas while dorsal ROIs are separated away from both early visual and lateral ROIs. These results provide evidence for distinct representational content along the two pathways.

Embodied cognition theories argue that recognition of observed actions relies on sensory-motor simulation and taps into motor representation necessary for executing those actions (Rizzolatti & Craighero, 2004; Pulvermüller, 2013). Motor, premotor, and parietal regions that contain mirror neurons in Macaque monkeys have been proposed as potential regions contributing to action recognition through motor simulation (Rizzolatti & Craighero, 2004). In our study we failed to find position tolerant represenations in prefrontal and premotor regions, and the representations in our parietal regions did not show correlation with the model based on subjective judgments on actions. These results suggest that the motor, premotor and parietal regions are unlikely to contribute to the subjective understanding of actions (Caramazza et al., 2014). Nevertheless, it is still possible that parietal regions would contribute to action observation during coupled motor interactions as well as motor imitations (Caspers et al., 2010) through analysis of body part movements. Our results suggest distinct roles for dorsal and lateral regions in action observation.

In summary, we found regions capable of representing highly abstract forms of observed human actions, namely the point light display video clips of actions across changes in position. These regions located in lateral occipito-temporal and parietal cortices contained distinct representational content. The lateral regions reflected more strongly the human subjective knowledge about objects, while the dorsal regions reflected only the objective bodily movements. These results suggest the existence of two distinct networks that contain abstract representations of human actions likely serving different purposes in the visual processing of actions.

